# PaxDB 5.0: curated protein quantification data suggests adaptive proteome changes

**DOI:** 10.1101/2023.04.16.536357

**Authors:** Qingyao Huang, Damian Szklarczyk, Mingcong Wang, Milan Simonovic, Christian von Mering

**Affiliations:** Swiss Institute of Bioinformatics and Department of Molecular Life Sciences, University of Zurich, Winterthurerstrasse 190, Zurich, 8057, Zurich, Switzerland

**Keywords:** protein abundance, proteomics, mass spectrometry, proteome evolution, database

## Abstract

The “Protein Abundances Across Organisms” database (PaxDB) is an integrative meta-resource dedicated to protein abundance levels, in tissue-specific or whole-organism proteomes. PaxDB focuses on computing best-estimate abundances for proteins in normal/healthy contexts, and expresses abundance values for each protein in “parts per million” (ppm) in relation to all other protein molecules in the cell. The uniform data re-processing, quality scoring, and integrated orthology relations have made PaxDB one of the preferred tools for comparisons between individual datasets, tissues or organisms. In describing the latest version 5.0 of PaxDB, we particularly emphasise the data integration from various types of raw data, and how we expanded the number of organisms and tissue groups as well as the proteome coverage. The current collection of PaxDB includes 831 original datasets from 170 species, including 22 Archaea, 81 Bacteria and 67 Eukaryota. Apart from detailing the data update, we also show a comparative analysis of the human proteome subset of PaxDB against the two most widely-used human proteome data resources: Human Protein Atlas (HPA) and Genotype-Tissue Expression (GTEx). Lastly, we present a use case of PaxDB, showing how protein abundance data can be used to study the evolution of relative amino acid usage in Fungi.

## 1 Introduction

Biological processes are regulated at multiple levels. Although many cellular changes are clearly detectable already at the transcriptome level, it is the protein level that most accurately reflects the cellular state, since proteins act as the direct executors of biological functions. Apart from a proteins’ expression level, further regulatory potentials lie in its post-translational modifications, sub-cellular localizations and biological contexts. In a complex multicellular organism with a system of coordinated organs, protein expression patterns largely conform to the specific requirements and activity of the tissue or organ. Furthermore, protein expression profiles can differentiate between healthy and disease states, providing important markers and targets for diagnosis and treatment.

Thus, systematic measurements of protein expression levels facilitate both the understanding of fundamental biological processes and the design of new therapeutic strategies. Proteomics data collections have seen exponential growth in the last decade. Along with the data growth, analytical instrumentation and data processing methodologies for quantitative proteomics have rapidly progressed. Mass-spectrometry based measurements provide the bulk of protein quantifications, with multiple workflows and modalities from stable isotope labelling to label-free quantification, from targeted, data-dependent acquisition (DDA; or shotgun) to data-independent acquisition (DIA) modes, and involving multiple ion trap technologies - time-of-flight (TOF) [1], linear quadrupole ion trap [2], orbitrap[3] - in terms of instrument configuration. For downstream data processing, a number of quantitification software packages evolved with multiple pipelines to tackle different challenges in each experiment set-up, with the most prominant ones being MaxQuant (MQ) [4] and Proteome Discoverer (PD; commercial). A plethora of file formats are produced; input and output data at several levels of processed information are recorded in various forms with overlapping information content [5]. Despite efforts to create a unified data standard with mzML [6] and mzTab [7], the legacy of viable file formats continues to create challenges for integrating and standardizing the existing data.

The PaxDB database (<u>P</u>rotein <u>A</u>bundances A<u>cross</u> Organisms) is an integrative meta-resource dedicated to absolute abundance levels in whole-organism or tissue-specific proteomes[8, 9]. PaxDB focuses on creating a consensus view on normal/healthy proteomes, and expresses abundance values in “parts per million” (ppm) in relation to all other protein molecules in the sample. Since the last PaxDB update, the proteomics community has grown continuously: roughly 1000 projects per month are submitted to ProteomeXchange, the largest centralized platform for MS-derived primary data submission [10], involving PeptideAtlas [11], PRIDE [12], iProx [13] and JPOST [14] among others. For the latest version 5.0 of PaxDB, we have further improved data integration by extending the types of raw data imported from the various repositories, and by expanding the number of organisms and tissue groups as well as the proteome depth of previously covered organisms.

Using earlier versions of PaxDB as a reference, scientists have already modeled fundamental biological processes[15–19], formed hypothesis about stoichiometry in complexes [20, 21], studied tissue-specific functionalities [21–23] and verified new MS techniques and methodology [24, 25]. Indeed, the overall protein abundance landscape is likely reflecting a fundamental, cross-species, structural and functional equilibrium [8]. Proteins at the high end of the abundance distributions are particularly informative for evolutionary studies: their synthesis brings a significant cost to the organism, and they are observed to be coded more compactly, to have fewer introns and to be subject to heavier codon optimizations [26–29]. As the biosynthetic energy costs of the various amino acids differ by as much as 7-fold, energetic effects - but also nutrient and element availability - shape the general direction of amino acid evolution[30, 31]. [32]. Episodes of nitrogen (N) limitation likely have lead plants to reduce the overall nitrogen presence in their proteome as compared with animal proteomes [33]. Iron (Fe) limitation prompted most marine organisms to develop an iron-free version of ferredoxin, flavodoxin, as an electron transfer agent in their biochemical reactions [34, 35], and in extreme environments, a clade of *Procholorococcus* permanently lost ferredoxin in addition to losing 10% Fe-containing proteins [36]. Sulfur (S) is somewhat less studied. It is present in only two amino acids’ side chains, cysteine and methionine. Nevertheless, it has been shown that the effect of a single amino acid substitution involving sulfur is visible to selection in more than half of the proteome in a yeast model [37]. While previous studies reached their conclusion through observations in a limited number of species and proteomics datasets, the PaxDB resource has the advantage of its large collection of protein abundance data with associated orthology relationships. Here, we use this data to present evidence of a strong and wide-spread sulfur avoidance at evolutionary timescales, in an entire clade of Fungi species.

## 2 Experimental procedures

### 2.1 PaxDB data

All data contained in PaxDB are derived from public repositories, open-access publications, or publicly accessible data supplements.

#### 2.1.1 Abundance data inherited from PaxDB v4

Protein records in PaxDB are generally based on the same genome versions and identifier namespaces as those in the STRING database [38], including one-to-one mappings to Uniprot IDs. PaxDB v4 corresponds to STRING v10.0, whereas the updated PaxDB v5.0 corresponds to STRING v11.5 [38]. Wherever genome annotations and protein sequences change between major PaxDB releases, the corresponding protein records are re-mapped to the newest annotation.

For datasets quantified by spectral counting, a re-computation is performed on the updated species’ complete proteome. For datasets consisting of protein identifiers and abundances, the protein names are re-mapped to the latest name-spaces using the identifier collections maintained by STRING, which provide identifier mappings for a total of 278 identifier systems.

#### 2.1.2 Dataset collection

Since 2014, the ProteomeXchange Consortium [10, 39] has become a centralized platform for MS proteomics data sharing. The selection of projects to be imported into PaxDB is based on project metadata and text mining of publications. The metadata of the projects were downloaded through the ProteomeXchange API, including project ID, species, year, keywords, list of files with extension, among other information. A full-text search was performed on all PubMedCentral publications and those containing identifiers starting with PXD or RPXD were retrieved. Combining the project metadata and publication information, the relevant projects were manually selected with priority for highly-cited publications and organisms not yet included in PaxDB. Keywords involving disease conditions, subcellular compartments, exclusively post-translational modification or protein identification, virus, secretome, interactome and metabolome were excluded.

During the project selection, the abundance data were downloaded from the supplementary tables in 98 publications.

An additional 421 project IDs were selected and their files were downloaded from ProteomeXchange. The file extensions which were further processed included: csv, xls/x, mzid, mgf, mzML, mzTab, msf and txt.

#### 2.1.3 Protein-centric data processing

From publication supplements, abundance reports already aggregated to the protein-level were directly mapped to STRING IDs using global alias files. To recover any unmapped entries, additional steps were taken. In particular, protein IDs in the International Protein Index namespace (IPI; closed with its last update in September 2011) system were converted to UniProt ID with the mapping file (last release 2014-01) from UniProt Archive (UniParc). The converted Uniprot IDs as well as other unmapped IDs were searched through the NCBI E-utilities [40] to fetch their protein sequences. These sequences were blasted against the updated species proteomes and mapped via reciprocal besthit matching (requiring a minimum 90% sequence identity in both directions). For datasets reporting protein abundances in the form of ortholog IDs from a closely related species (because mass spectrometry libraries and databases were available there), protein records were also blasted against the original species’ proteome and mapped via reciprocal best-hit matching.

In bulk-download files from ProteomeXchange, most protein-centric data are in the MaxQuant output format “proteinGroup.txt” [4]. Any “CON “ (contaminant) or “REV_ “ (reversed sequence) entries were excluded from proteinGroups files, while all other protein names were mapped as described above. The intensity value for each protein entry was extracted and the molecular weight and the theoretically observable peptides were calculated from the protein sequence. Then, the intensity-based absolute quantification (iBAQ) was calculated, using Method 4 described in [41].

#### 2.1.4 Peptide-centric data processing

Wherever available, peptide-centric data were preferred over protein-centric data, and peptide-intensity data were preferred over peptide-count data, for all downstream data processing. From msf files, target and decoy peptides were separated. The FDR score threshold was set to 0.01 to filter the valid target peptides. The peptide intensities were extracted for further processing. All imported peptide data were then further processed with the pipeline described in [42].

#### 2.1.5 Data quality control and dataset integration

Since interacting pairs of proteins tend to be expressed at broadly similar abundance ranges, global protein-protein interaction information can be used to derive an estimate of data quality for each dataset [8]. For version 5.0 of PaxDB, all interacting protein pairs are retrieved from STRING v11.5 [38]. For each dataset to be imported into PaxDB, the absolute log abundance ratios of interacting protein pairs are computed and the median is taken. Then, the same operation is executed 500 times for the same dataset but with shuffled protein labels. The z-score of the observed median against the distribution of medians after label shuffling is termed “interaction z-score”, with a larger value indicative of better overall data quality.

In case more than one independent dataset is available for a given organism or tissue group, an “integrated” dataset is generated by weighted averaging. The estimation of the weights is iterative: the datasets are sorted by their interaction z-scores; the best-scoring dataset receives a weight of 100%. Then, proceeding from second-best to the worst-scoring dataset, each new dataset is merged to the previously integrated datasets in ten differently-weighted attempts, and the weight of the best-scoring attempt is chosen.

#### 2.1.6 Metadata standards

Each dataset’s metadata, such as the organism name, taxonomy identifier (NCBI taxonomy [43]), tissue, tissue ontology ID, publication/source, PubMed ID and quantification method (free text) are collected and made available via the website as well as the download files. Wherever possible, tissues are encoded with one of the following ontology systems: Uber-anatomy ontology (UBERON) [44], Plant Ontology (PO) [45], Cell Line Ontology (CLO) [46], Cell Ontology (CL) [47] and The BRENDA Tissue Ontology (BTO) [48].

### 2.2 Human proteome comparisons

#### 2.2.1 Data collection

The Human Protein Atlas (HPA) normal tissue data based on version 21.1 and Ensembl version 103.38 was downloaded. 23 tissues were subsequently mapped to their corresponding PaxDB tissue categories with identical labels for all but two tissues (“heart muscle”, “saliva gland” in HPA and HEART, SALIVA SECRETING GLAND in PaxDB, respectively).

GTEx proteome data were collected from Supplementary Table S2 (protein level) from [49]. For each organ group, the GTEx consortium had quantified between two and eleven selected proteome samples per organ by mass spectrometry. The GTEx organ information of these samples was extracted and mapped, resulting in 13 PaxDB-matched organ groups. For the transcript expression data, gene-level TPM values were downloaded from the GTEx portal from Analysis V8 (dbGaP Accession phs000424.v8.p2), involving 17384 samples per organ. 15 organ groups were matched to PaxDB. For each organ group, the global means per gene across all samples was used.

#### 2.2.2 Data analysis

The HPA normal-tissue protein data was first filtered by the “Reliability” parameter: proteins in the “Uncertain” category were removed. The four protein levels: “Not detected”, “Low”, “Medium” and “High” were then used for the comparison. A one-way ANOVA was performed for each tissue with PaxDB abundance ppm values, using HPA protein level as groups.

The PaxDB human proteome data (excluding GTEx experiments) were compared against GTEx RNA and protein datasets. A tissue-specific z-score was calculated for each gene and each tissue in PaxDB integrated data and GTEx data to represent relative protein expression [49]. The Pearson correlation was calculated for z-scores between the tissue datasets. The Pearson’s correlation coefficients of organ groups from both paxDB and GTEx were hierarchically clustered using average linkage to generate a heatmap.

### 2.3 Proteome Evolution in Fungi

#### 2.3.1 Orthology and protein domain matching

179 *Fungi* species as well as five reference species from other Eukaryotic clades (*Homo sapiens, Drosophila melanogaster, Caenorhabditis elegans, Plasmodium falciparum and Dictyostelium discoideum*) were included in the study. The “Simple Modular Architecture Research Tool” (SMART) [50] was used to assign annotated domains to all encoded proteins in their genomes. Orthology relations between the genes in these species were retrieved from EggNOG version 5.0 [51]. For all matched domain pairs between a reference species and a given Fungi species of interest, pairwise global sequence alignments were performed with EMBOSS-needleall [52] using the Needleman-Wunsch alignment algorithm [53]. The domain pairs were filtered for at least 40% sequenceidentity, and only the best-scoring alignment pair was considered in case multiple domains of the same type were annotated for any of the two orthologous sequences.

#### 2.3.2 Amino acid usage ratios between Fungi and reference proteomes

For each orthologous protein pair, the amino acid usage was counted within the aligned domain sections, and a ratio was computed. Orthologous pairs were only considered if both proteins had at least one count each, of the amino acid of interest. In cases where the orthology relation was complex (i.e. where paralogous proteins are annotated in one or both of the organisms), the orthologous group was also not considered. For each species pair (Fungi species vs. reference species), the protein abundance values were taken from the orthologous protein in *S.cerevisiae*, using its WHOLE ORGANISM integrated data.

A Spearman’s rank correlation was then calculated between the amino acid usage ratios and protein abundances. For the visualizations in Fig. **??**, the data points were binned in six equally-sized groups for the violin plots, and a linear regression was fit over all data points.

A taxonomy tree visualization was created for the 179 Fungi species according to the NCBI taxonomy database [43], using the interactive Tree Of Life (iTOL) [54] online tool. A heatmap was then added to the tree, visualizing the Spearman’s r values for each comparison between a Fungi species and a reference species.

## 3. Results

### 3.1 Data update

The current PaxDB version 5.0 has nearly doubled its data content, both with respect to the number of datasets, as well as the number of organisms covered (Fig. 1 A). The PaxDB data integration pipeline involves a keyword-based discovery search for suitable projects and/or publications, and automatic data processing for multiple input formats downloaded from repositories. 184 of the 322 newly imported datasets were derived from open-access publications’ supplementary files (“Curated”) and 133 were from data repositories (“Repository”). From a data processing point of view, 209 of the datasets were passed through our protein-centric import pipeline, which involves protein name mapping and where necessary also mapping via protein sequence comparison. In addition, 113 datasets were passed through the peptide-based import pipeline. 25 of the latter were processed from msf files, and the rest from plain text inputs (Fig. 1 B).

**Fig. 1.**
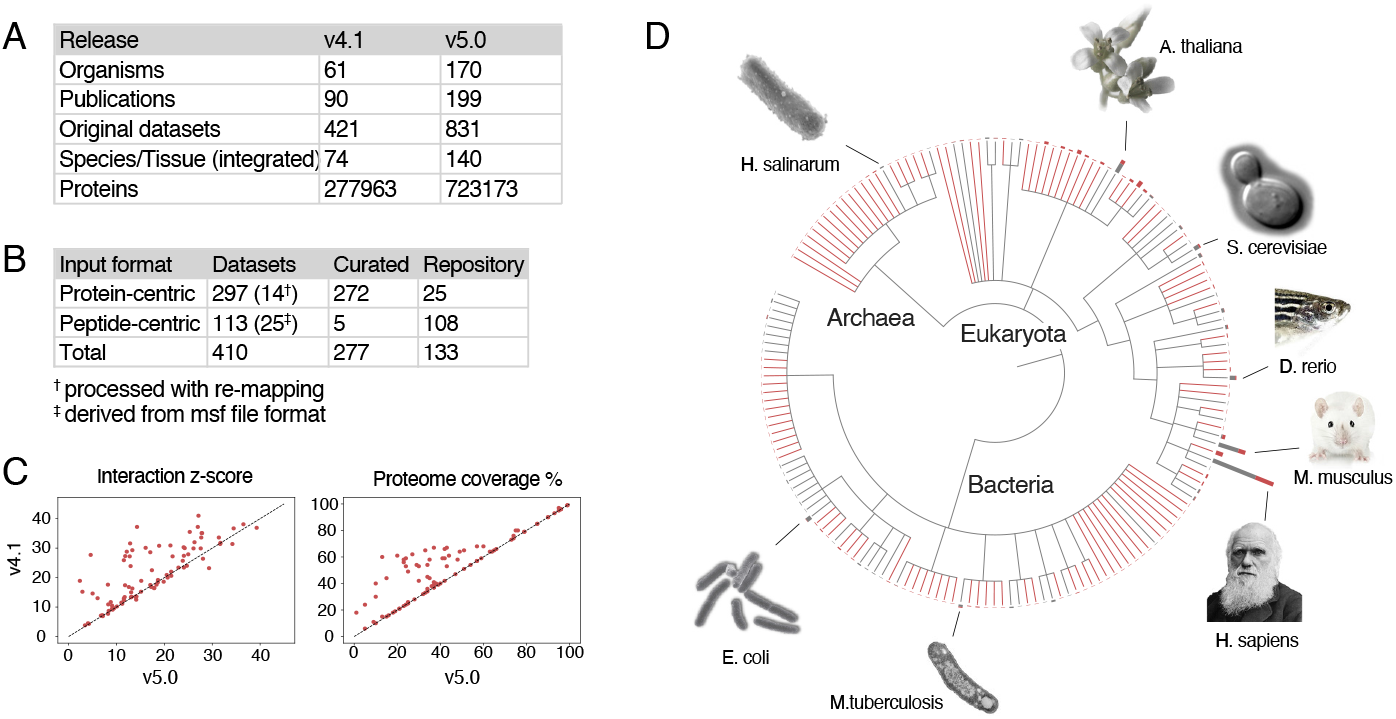
PaxDB v5.0 data overview. A: Comparison to the previous version (v4.1), in terms of number of organisms, publications, original and integrated datasets, peptide spectrum matches (PSMs) and proteins covered; B: Origins and input formats of newly acquired datasets in v5.0; C: Interaction consistency scores and proteome coverage for newly added as well as existing datasets. D: PaxDB 5.0 contains 170 species spanning 3 domains of life. Gray lines represent 61 species already existing in v4.1 and red lines represent 109 newly added species in v5.0. The associated bar plots indicate the number of datasets per species (Gray: existing; Red: new datasets).

Since genome annotations and name-spaces continually evolve, the datasets from version 4 of PaxDB also needed to be re-mapped and/or re-quantified. The datasets’ quality scores (i.e. their interaction consistency, see *Procedures* 2.1.5 above) were found to be mostly improved after the version update, when using the same STRING protein network as reference. For the “integrated” datasets (i.e. the best-estimate weighted combinations per species or tissue group) the interaction z-scores and proteome coverage nearly always increased when new datasets were added (see Fig. 1 C for a comparison of for 93 integrated datasets with their counterparts in version 4). For existing groups to which no new datasets were added, the scores stayed approximately the same. “New in v5.0” groups denote groups where only one original dataset existed in v4, with additional datasets in v5 now enabling an integration. All except one new group had similar or higher scores than before.

The 170 species in version 5.0 of PaxDB span the Archaea, Eukaryota and Bacteria domains (Fig. 1 D). The number of Archaea species increased from 1 to 22. While more Bacteria species are included, the Eukaryota still encompass the largest number of datasets, especially due to diverse tissue-differentiated measurements. The top 3 species in terms of the number of datasets are *H.sapiens* (249), *M.musculus* (106) and *A.thaliana* (59). *E.coli, H.sapiens* and *S.cerevisiae* are the top 3 species in terms of maximal proteome coverage, followed up by *B. subtilis, P.falciparum, D.melanogaster, B.burgdorferi, H.pylori* 26695, *M.musculus* and *S.pombe* which all had over 90% maximal proteome coverage.

### 3.2 Comparison to other human proteome resources

To independently assess the validity of PaxDB data, it was compared to two comprehensive gene expression resources focusing on *Homo sapiens*: the Genotype-Tissue Expression (GTEx) project as well as the Human Protein Atlas (HPA), both of which contain tissue-specific protein expression data.

#### 3.2.1 Genotype-Tissue Expression (GTEx)

Both RNA-level and protein-level data from GTEx samples were used. The tissue specificity z-score was calculated per gene to represent a gene-level signature per organ group for both GTEx and paxDB. The “signatures” from both resources were clustered by their pairwise Euclidean distance and a heatmap was colored by the Spearman correlation coefficients. On protein level, the matched organ group pairs between paxDB and GTEx which also clustered together included adrenal gland, brain cerebral cortex, lung, liver, pancreas, spleen, thyroid gland, and testis (8 out of 13; Fig. 2A On RNA level, kidney, adrenal gland, colon, brain cerebral cortex, testis, esophagus (partially), stomach, spleen, fallopian tube were clustered together (9 out of 15; Fig. S1). T tests for in-group and out-group Euclidean distances showed significance for RNA (p-value 2.28 × 10^− 2^) and protein (p-value 3.64 × 10^− 5^). Some organ groups contain multiple tissue lineages, e.g. stomach, esophagus, colon have epithelial and muscle layers. Their sampling proportions could contribute to the difference in the expression landscape. Other organs were previously reported to be relatively indiscriminate, e.g. prostate [49]. In addition, the difference could also be attributed to lower proteome coverage, e.g. the skin dataset from paxDB (excluding GTEx) covered only 23% proteome, which likely reduced its tissue-distinguishing potential.

**Fig. 2.**
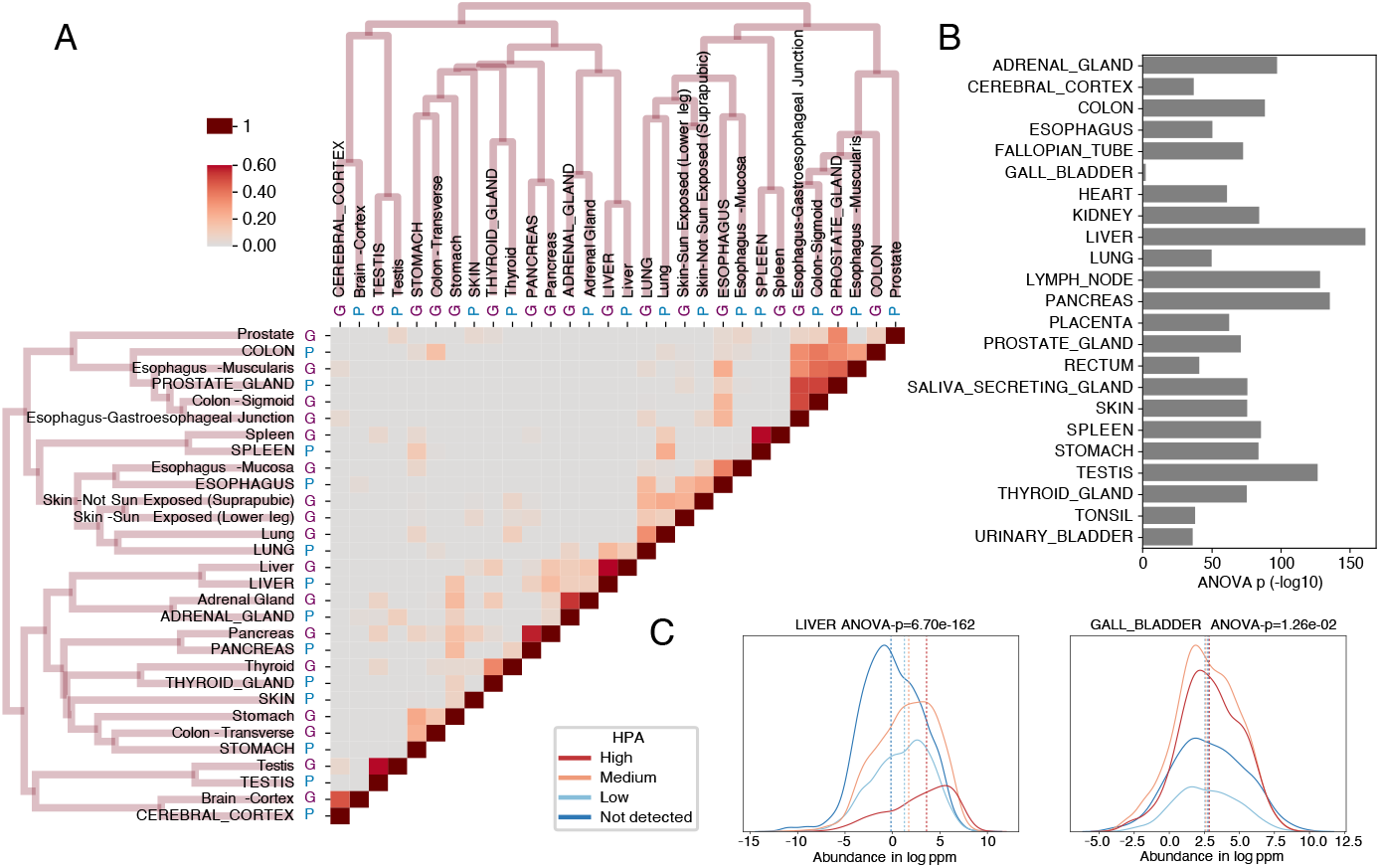
Human proteome quantifications across distinct database resources. PaxDB quantifications are compared to Human Protein Atlas and GTEx data. A: Pearson’s correlation for tissue specific abundances in PaxDB against GTEx protein with clustering dendrogram, with PaxDB tissues marked with P and GTEx tissues with G. B: The PaxDB protein abundance distributions of human proteins, grouped by broad “expression level” categories from HPA, in liver and gallbladder. C: Significance of abundance separation by HPA group label, per human tissue type.

#### 3.2.2 Human Protein Atlas (HPA)

While PaxDB data is almost entirely derived from MS data, the HPA normal tissue datasets approximates the protein expression profiles with antibody-based tissue micro-arrays. PaxDB computes protein abundance in continuous ppm values, while HPA reports protein expression in four levels, from “high”, through “medium” and “low”, to “undetected”. The different characteristics of either technology may result in systematic biases in the results, but a global abundance trend is expected to be observable in both. In Fig. 2C, the overlapping proteins were grouped by HPA labels and the distribution of protein abundance according to PaxDB with a vertical median line was shown. In kidney, the median separation from “high” to “not detected” was evident with ANOVA p-val at 8.6 × 10^− 88^; whereas in skin, only the “high” group had elevated abundance from the other groups, with ANOVA p-val at 5.27 × 10^− 7^. Across the 23 tissues, the ANOVA p-val varied between\ 1.69 × 10^− 2^ in GALL BLADDER, and 1.19 × 10^− 129^ in LYMPH NODE (Fig. 2 D, red crosses). The tissue-shuffled ANOVA p-value medians ranged between 8.62 × 10^− 5^ (SKIN) and 4.22 × 10^− 86^ (TESTIS), while the p-values with the correct tissue label generally showed more significant difference between group labels. In 13/23 tissues, the z-scores of the real p-value in the shuffled distribution were significant (*α* = 0.05; Fig. 2 B upper panel), indicating high tissue-specificity in the proteome data.

### 3.3 Use case: evolution of sulfur-containing amino acids in Fungi

The relative frequencies of amino acids in the overall proteome can change at evolutionary timescales [55] and they are known to differ across organisms in response to mutational and environmental processes [56–58]. While inspecting datasets in PaxDB, we noticed that sulfur-containing amino acids (cysteine and methionine) seemed to be markedly under-represented in the proteomes of certain Fungi. On its own, this observation is difficult to interpret: it could be the consequence of distinct functional compositions of these proteomes (e.g. unusual fractions of secreted proteins), or the result of genome-wide mutational biases, or it could suggest an adaptive response. To narrow down potential causes, we compared Fungi proteomes to a varied set of other eukaryotic proteomes, and restricted the comparisons to strictly one-to-one orthologous gene pairs 3 In addition, we further restricted the analysis to functionally and structurally equivalent parts within these orthologs (i.e., aligned protein domains); doing so should largely cancel out effects caused by differences in overall proteome functions. Then, to distinguish genome-wide mutational effects (e.g. G/C content differences) from potentially adaptive effects, we further stratified proteins by their absolute abundance levels - adaptive changes in response to sulfur limitations should be visible particularly in highly-expressed proteins.

We systematically conducted this analysis on the proteomes of 179 Fungi species, for which the encoded proteomes, protein domain compositions and orthology relations have been established [38, 50, 51]. We compared their sulfur usage against five representative eukaryotes from other clades, selected for high proteome coverage in PaxDB. The latter included the human, two animal model organisms (*C.elegans* and *D.melanogaster*) and two unicellular eukaryotes (*P.falciparum* and *D.discoideum*).

When comparing the ratios of sulfur-containing amino acids across orthologous gene pairs, we observed for the majority of proteins expressed at low to medium levels, that the overall usage of sulfur was roughly similar (i.e., the median ratio centered on 1.0, see Fig. 3). However, remarkably, this ratio dropped below 1.0 for more strongly expressed, abundant proteins. This trend is highly significant, and is observed independently both for cysteine as well as for methionine. It also made no difference whether protein abundance levels were taken from one or the other of the two organisms (not shown); the yeast (*S.cerevisiae*) was taken as a reference for protein abundances for all subsequent analysis because of its highest quality and coverage in paxDB, within the Fungi clade.

**Fig. 3.**
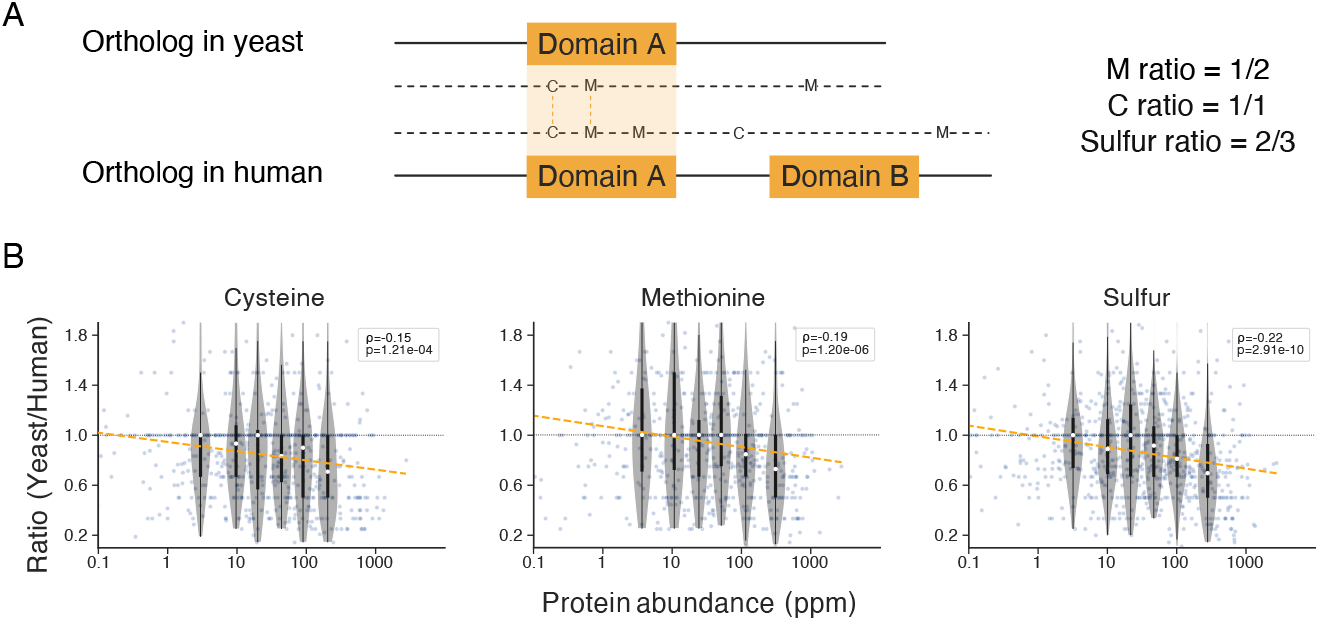
Comparing sulfur usage in orthologous protein pairs. A: To avoid potentially confounding effects of evolutionary changes in protein function, only equivalent and aligneable protein domains within strict 1:to:1 orthologs are considered. B: Relations between protein abundance and the sulfur-usage ratio yeast/human. Each data point corresponds to one orthologous pair of proteins; the abundance values on the x-axis are taken from the yeast protein. Violin plots indicate median and percentiles 25 and 75, for six equally sized bins. A linear regression was fit for the regression line. Spearman’s r and p-value were separately derived.

Assessing the strength of this effect across all 179 Fungi, a differential sulfur depletion pattern was observed (Fig. 4) Separate heatmaps for cysteine and methionine shows coordinated regulation patterns (Pearson’s r ranges between 0.52 and 0.74 against five reference species; Fig. S2 and S3). While the majority of Fungi species show at least some degree of sulfur avoidance against the five reference species, the reduction was the strongest in the *Saccharomycotina* order (C in Fig. 4), containing baker’s yeast as well as most other unicellular Fungi species (i.e., yeasts). Assuming that the observed sulfur avoidance in the Saccharomycotina might be adaptive, i.e. perhaps related to recurring episodes of sulfur limitation, a multicellular lifestyle would have been advantageous as it could provide mobility to escape the limiting environments. In the Fungi kingdom, the marker for multicellularity is the development of hyphae, long tubular substrate-seeking extensions, which allow the organisms to survive and migrate away from nutrient-poor areas [59]. We asked whether other unicellular yeasts besides Saccharomycotina showed elevated sulfur avoidance. As the unicellularity in Fungi is not monophyletic, we marked the uni-/multicellularity of the species in Fig. 4 according to MycoBank [60]. Although organisms in the *Taphrinomycotina* subdivision (B in Fig. 4) and a few species in the *Basidiomycota* division (A in in Fig. 4) are also unicellular, their proteomes did not exhibit similar levels of sulfur reduction as the Saccharomycotina. We retrieved genome-wide median GC% from NCBI Genome for each fungi species. While with all species, it appeared to indicate a positive correlation between GC% and the sulfur ratio (Pearson’s r 0.38, p-val 2.3 × 10^− 7^), excluding the Saccharomycotina clade removed the correlation completely (Pearson’r 0.01, p-val 0.9). This indicates an associated effect of the lower GC content of Saccharomycotina clade. We further investigated the environment/host associations of the Saccharomycotina and the closely related Taphrinomycotina species. Habitat and/or lifestyle information for Fungi is not systematically available; we approximated it by the annotated sources of the first isolation of the type strains, as described in ATCC [61]; we also marked potential symbiontic lifestyles with “c” and parasites with “p” and free-living lifestyle with “f”. For Taphrinomycotina (showing little sulfur avoidance), two out of six species were parasitic, while for Saccharomycotina (strong sulfur reduction), 11 out of 33 species were symbiont, six of which were parasitic. Overall, a clear association of sulfur avoidance with free-living or parasitic lifestyles was not observed. As the environments of the isolated type strain cannot fully represent the major habitat of the species, the underlying cause of the clade-wide sulfur avoidance was not established.

**Fig. 4.**
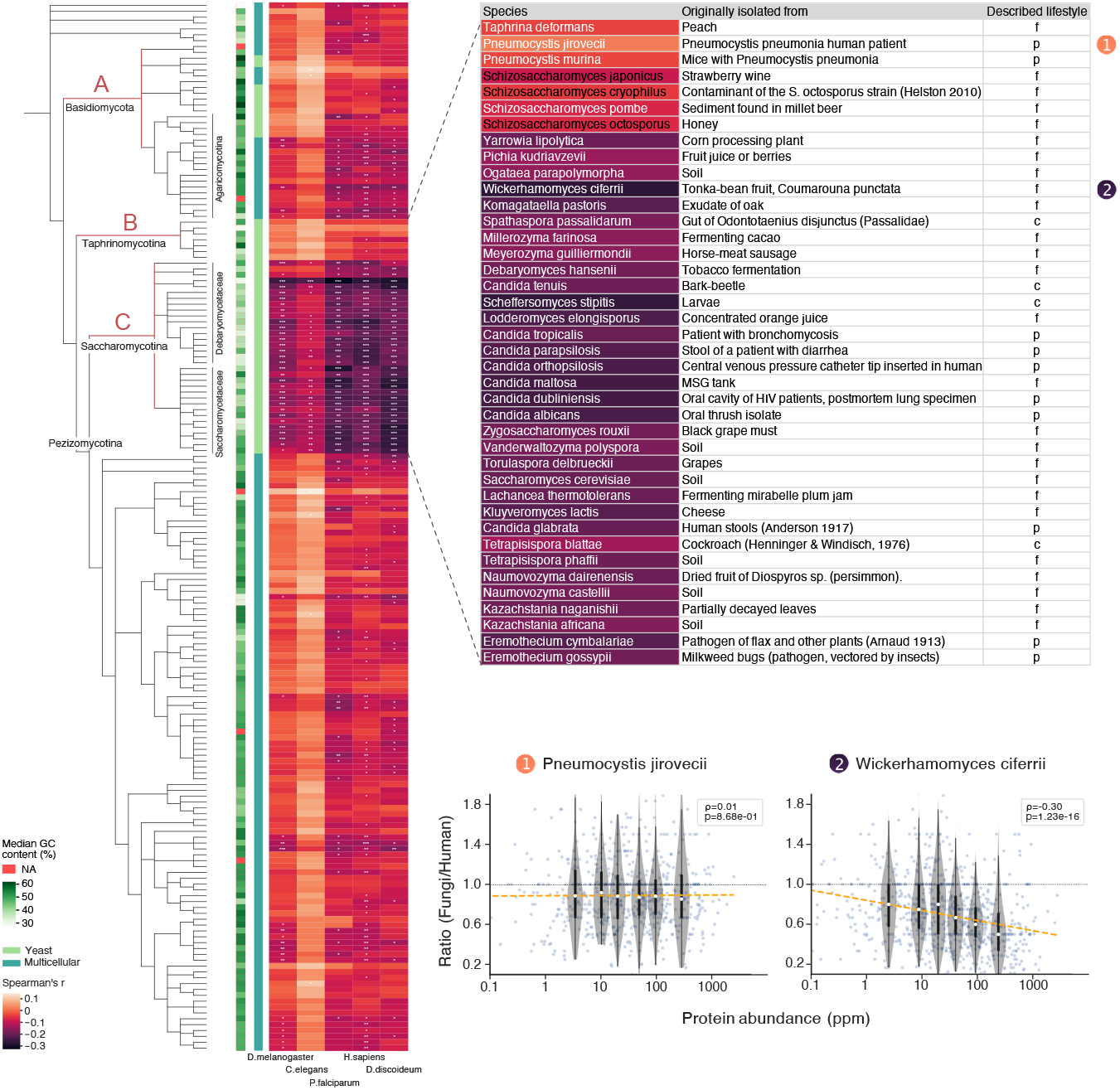
Patterns of reduced sulfur content in Fungi proteomes. The proteomes of 179 Fungi (rows) are compared to the proteomes of five reference organisms from other Eukaryotic clades (columns). Each tile in the heatmap indicates the strength of the negative correlation (Spearman’s r) between protein abundance and the sulfur-usage ratio Fungi/Reference. Asterisks indicate the significance (p-value) of the correlations: *: *<* 0.01, **: *<* 0.001, ***: *<* 0.00001. Each Fungi’s genomic GC-content and multicellularitly status are color-labeled on the left of the heatmap. The Fungi are arranged according to their taxonomic annotations at NCBI [43]. The Taphrinomycotina (B) and Saccharomycotina (C) clade are shown expanded to the right, together with the heatmap colors of the Fungi/human comparisons; the inset includes information on the environment from which the original type strains were collected. Two Fungi are highlighted (arrows); their detailed correlation data are shown in the inset below, similar to Fig. 3.

## 4 Discussion

This update of PaxDb v5.0 reports a nearly three-fold increase in the number of species covered, and a two-fold increase in the number of original datasets and publications. Decreased evolutionary distances between the species will enable higher-resolution cross-species comparisons. Likely due to general improvements of the genome/proteome reference annotations, the re-mapping of older datasets from the previous version of PaxDb mostly resulted in dataset quality improvements.

Using two independent human-centered data collections - Human Proteome Atlas (HPA) and Genotype-Tissue Expression (GTEx), we verified the overall data quality of PaxDb, in terms of quantitive agreement and dataset tissue matching. We compared the PaxDB integrated tissue-level protein abundances with matched GTEx RNA and protein abundances. While both showed strong correlation between the matched labels, the protein-level comparison contained fewer indiscriminate correlations and more in-group correlations than the RNA-level. Comparisons with antibody-based protein abundance estimates from HPA reached qualitatively similar conclusions. Using proteinprotein interaction data as another, independent arbiter of data quality, we show that the PaxDB integration of multiple lower-coverage, lower-quality datasets enhances the data quality and provides a boost to overall proteome coverage.

By integrating PaxDB data with sequence analysis of orthologous protein pairs, we discovered an apparent, strong selection pressure to reduce sulfur usage in abundantly expressed proteins, in a particular clade of single-celled Fungi. One of the conceiveable selection pressures causing this effect would be a recurring sulfur limitation in the environment. Experimentally induced sulfur depletion was shown to trigger an alternative proteome state, resulting in 30% reduction in sulfur usage in Fungi [62] and 45% reduction in a green alga, *Chlamydomonas reinhardtii* [63]. Besides transient responses to sulfur limitation in environment, sulfur limitation may also have resulted in adaptive changes in the genome. Baudouin-Cornu et al. showed, for example, that sulfur assimilatory enzymes in yeast and E.coli are themselves encoded using remarkably little cysteine and methionine [64]. Comparison of Cyanobacteria strains isolated from sulfur-rich and sulfur-poor environment showed adaptive eradication of C and M in phycobilisome, the light-harvesting proteins and the major cellular component, in response to sulfur depletion [65]. Another possible selective pressure against sulfur usage relates to oxidative toxicity. Unwanted disulfide bonds may be formed under oxidative stress, impacting protein folding and activity [66]. If the organisms were exposed to oxidative stresses through their evolution, it could explain the reduction of cysteine (but not methionine) in their protein sequences.

However, why the *Saccharomycotina* in particular would show a reduced use of sulfur in their proteome remains unclear. The habitat ranges and ecological strategies of many Fungi are described only anecdotally, and even less is known about any present or past episodes of sulfur limitations. Nevertheless, Fungi are known to be able to assimilate sulfur from a number of sources, both of biotic and abiotic origin [67]. Perhaps this diversity of assimilatory toolkits is a sign for past exposure to sulfur limitation. Future growth in the availability of genome sequences will allow this phenotype to be mapped with ever increasing resolution.

## 5 Abbreviations

FDR: False discovery rate;
UBERON: Uber-anatomy ontology;
PO: Plant Ontology;
CLO: Cell Line Ontology;
CL: Cell Ontology;
BTO: The BRENDA Tissue Ontology;
HPA: Human Protein Atlas;
GTEx: Genotype-Tissue Expression;

## Supplementary information

This article is accompanied by Supplementary Information (Figures S1, S2). The proteome abundance data are freely available at https://pax-db.org/.

## Acknowledgments

We would like to thank Dr. David Lyon for the documentation and project maintenance work, and current and former members of the von Mering group for valuable discussions and input. We also thank the Swiss Institute of Bioinformatics (SIB) and Swiss National Science foundation (SNSF) for the funding support.

## Notes

https://www.pax-db.org

### Competing Interest Statement

The authors have declared no competing interest.

