## Supplementary Material for "PaxDB 5.0: curated protein quantification data suggests adaptive proteome changes"

### Table of Content

Supplementary Figure 1: Pearson's correlation for tissue specific abundances in PaxDB against GTEx RNA expression data with clustering dendrogram, with PaxDB tissues marked with P and GTEx tissues with G.

Supplementary Figure 2: The proteomes of 179 Fungi (rows) are compared to the proteomes of five reference organisms from other Eukaryotic clades (columns) with respect to cysteine (left), methionine (middle) and sulfur.

Supplementary Figure 3: Correlations of cysteine and methionine ratio with abundance are verified with five species in a coordinated manner.

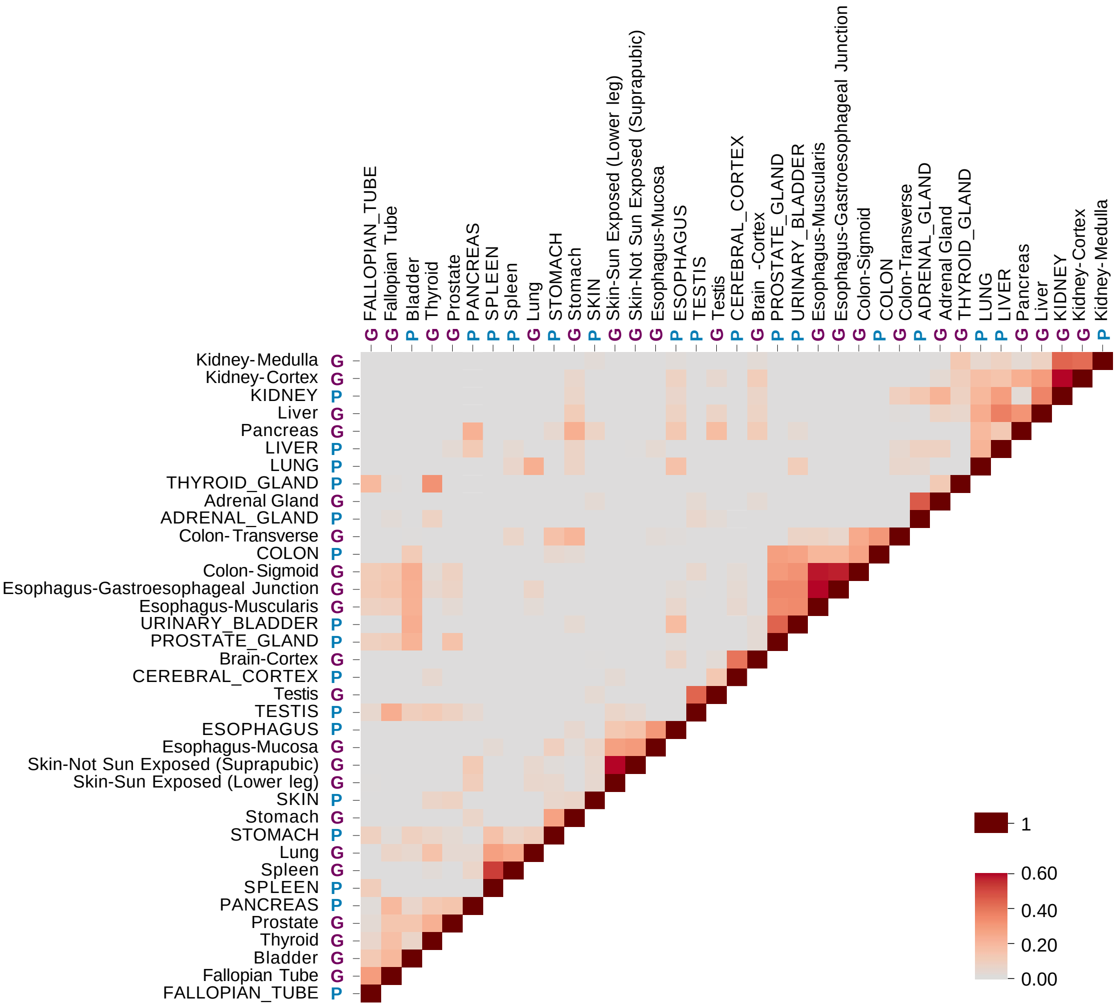


Supplementary Figure 1: Pearson's correlation for tissue specific abundances in PaxDB against GTEx RNA expression data with clustering dendrogram, with PaxDB tissues marked with P and GTEx tissues with G.

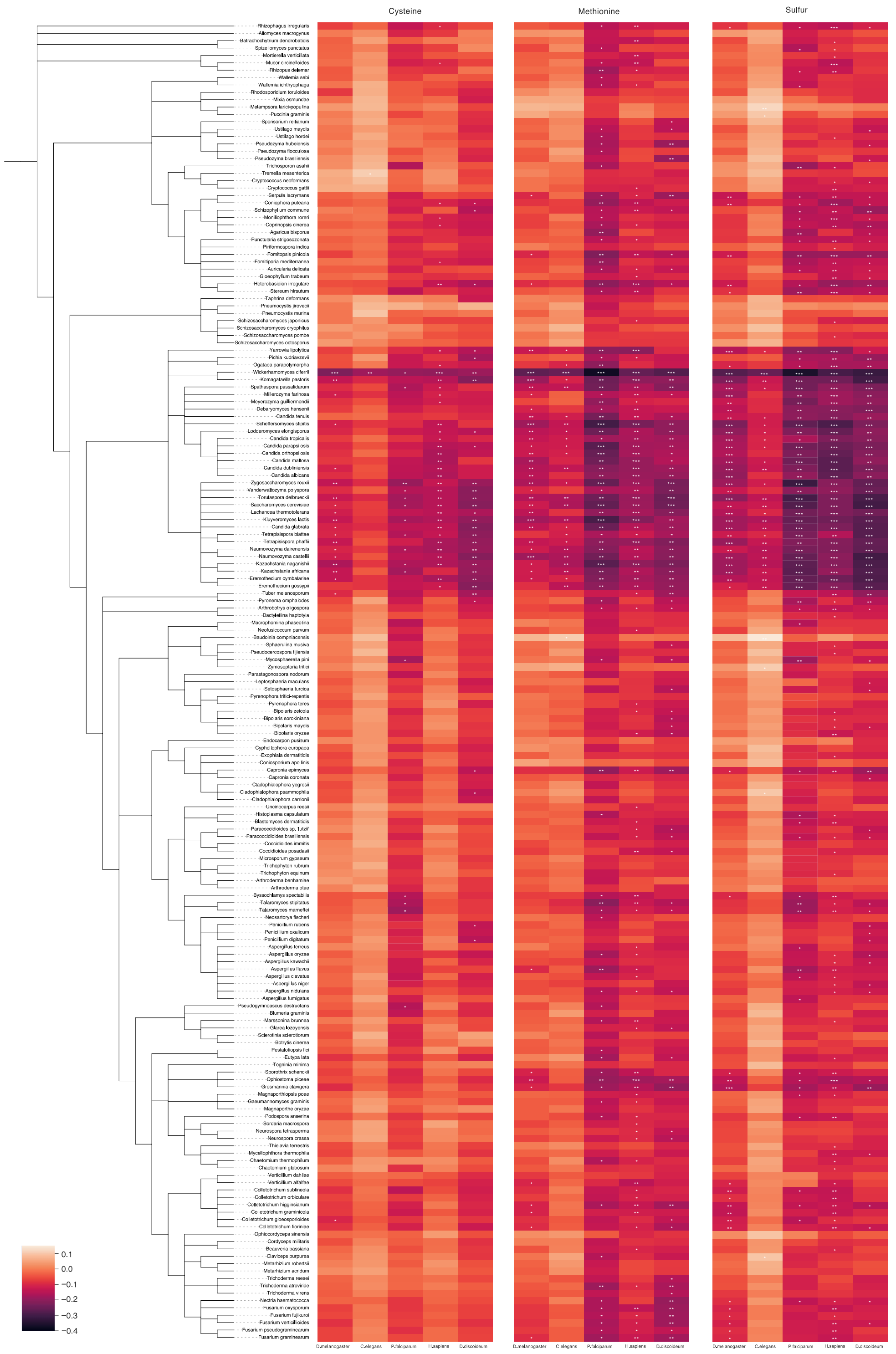


Supplementary Figure 2: The proteomes of 179 Fungi (rows) are compared to the proteomes of five reference organisms from other Eukaryotic clades (columns) with respect to cysteine (left), methionine (middle) and sulfur.
Each tile in the heatmap indicates the strength of the negative correlation (Spearman's r) between protein abundance and the sulfur-usage ratio Fungi/Reference. Asterisks indicate the significance (p-value) of the correlations: *: < 0.01, **: < 0.001, ***: < 0.00001.

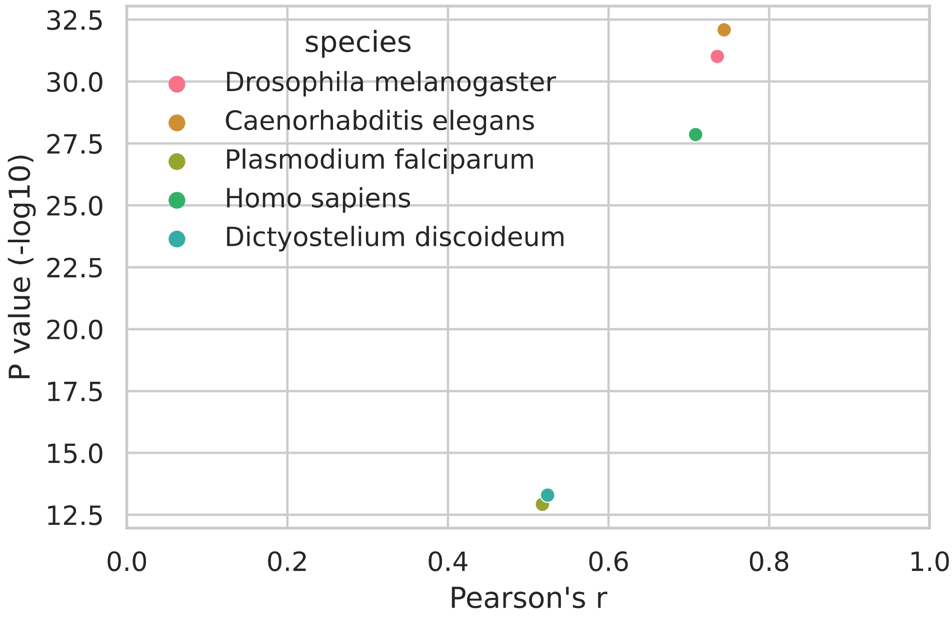


Supplementary Figure 3: Correlations of cysteine and methionine ratio with abundance are verified with five species in a coordinated manner.
The Pearson’s correlation test between cysteine and methionine for the Spearman’s *rho* across Fungi species shown in Fig. S2 indicates positive correlation (Pearson’s r ranges from 0.52 to 0.74 with significance (p-value ranges from $1.19 \times 10^{-13}$ to $8.17 \times 10^{-33}$).
